# Inferring the Evolutionary Reduction of Corm Lobation in *Isoëtes* using Bayesian Model-Averaged Ancestral State Reconstruction

**DOI:** 10.1101/212316

**Authors:** Forrest D. Freund, William A. Freyman, Carl J. Rothfels

## Abstract

Inferring the evolution of characters in *Isoëtes* has been problematic, as these plants are morphologically conservative and yet highly variable and homoplasious within that conserved base morphology. However, molecular phylogenies have given us a valuable tool to test hypothesis of character evolution within the genus. One such hypothesis is that extant *Isoëtes* have undergone a morphological reduction from larger arborescent lycophyte ancestors. In this work we examine the reduction in lobe numbers on the underground trunk, or corm, over evolutionary time. Using reversible-jump MCMC and Bayesian inference, our results support the hypothesis of a directional reduction in lobe number in *Isoëtes*, with the best-supported model of character evolution being one of irreversible reduction. Furthermore, the most probable ancestral corm lobe number of extant *Isoëtes* is three, and a reduction to two lobes has occurred at least six times. From our results, we can infer that corm lobation, like many other traits in *Isoëtes*, shows a degree of homoplasy, yet also shows ongoing evolutionary reduction.

## INTRODUCTION

Interpreting the morphological evolution of *Isoëtes* L. has troubled botanists for many years. These plants have a outwardly simple, highly conserved body plan, consisting of an apical rosette of linear sporophylls on top of a reduced corm-like trunk (Engelmann, 1882; Pfeiffer, 1922; Cox and Hickey, 1984; Hickey, 1986; Taylor and Hickey, 1992; Budke et al., 2005), yet are highly variable both within (Budke et al., 2005; Liu et al., 2006) and among closely related species (Cox and Hickey, 1984; Hickey, 1986; Taylor and Hickey, 1992; Romero and Real, 2005; Bagella et al., 2011). Even characters once thought to be useful in delimiting natural groups within the genus, such as habitat (Engelmann, 1882) or megaspore morphology (Pfeiffer, 1922), have been found to be labile (Cox and Hickey, 1984; Taylor and Hickey, 1992; Budke et al., 2005; Hickey, 2007; Bagella et al., 2011). And while some characters such as the glossopodium (the portion of the ligule internal to the leaf) have shown some potential (Sharma and Singh, 1984; Pant et al., 2000; Shaw and Hickey, 2005; Singh et al., 2010; Freund, 2016), actually examining and interpreting these structures requires considerable histological and computational effort, making them ill-suited for field identification. This absence of consistent, dependable characters creates a paradox: the lack of reliable traits impedes the inference of phylogenies or classifications in the genus, but without a phylogeny, examining character evolution is exceptionally difficult.

However, molecular phylogenetics has helped to vastly improve our understanding of *Isoëtes* (Hoot et al., 2004; 2006; Schuettpelz and Hoot, 2006; Larsén and Rydin, 2016), and has provided evidence for five major clades of *Isoëtes*: the Gondwanan Clade (Clade A), Laurasian Clade (Clade B), Italian Clade (Clade C), Austro-Asian clade (Clade D), and the New World or American Clade (Clade E; Hoot et al., 2006; Larsén and Rydin, 2016). This phylogeny has completely overturned the old morphological and ecological systems of classification, and has also provided a framework to begin earnestly studying character evolution in the genus. While characters such as spore ornamentation are highly homoplastic (Cox and Hickey, 1984; Larsén and Rydin, 2016), there are other changes that may be informative, such as serial reduction of the corm (Karrfalt and Eggert, 1977a; b; Pigg, 1992; Grauvogel-Stamm and Lugardon, 2001).

The corm of *Isoëtes* is quite different from the identically named structure of spermatophytes. In addition to possessing independently evolved secondary growth (Scott and Hill, 1900; Stokey, 1909; Pfeiffer, 1922; Stewart, 1947; Karrfalt and Eggert, 1977b), it has a unique capacity to penetrate into the substrate by laterally displacing the soil and pulling itself deeper through a combination of said secondary growth and development of new rootlets (Karrfalt, 1977). The corm of modern *Isoëtes* and the stigmarian appendages of extinct arborescent lycophytes are quite similar (Stewart, 1947; Karrfalt and Eggert, 1977a; b), to the point that they have been hypothesized to be homologous structures (Karrfalt and Eggert, 1977a; b; Jennings et al., 1983; Pigg, 1992).

Some fossils, such as *Protostigmaria* Jennings (Jennings et al., 1983) or *Nathorstiana* Richter (Taylor et al., 2011), have corm morphology comparable to that of modern *Isoëtes*, with only the size, elaboration, or arborescent habit of the plants separating them (Stewart, 1947; Karrfalt, 1984a; Pigg, 1992). Due to this similarity, it has been hypothesized that the morphology of modern *Isoëtes* is the result of reduction of the once arborescent isoëtalian body plan, and modern bilobate *Isoëtes* represent reduction from an ancestral trilobate corm (Stewart, 1947; Karrfalt and Eggert, 1977a; b; Jennings et al., 1983; Pigg, 1992).

These corm lobes are formed by the basal, root-producing meristems (Engelmann, 1882; Stokey, 1909; Osborn, 1922; Pfeiffer, 1922; Bhambie, 1963; Karrfalt and Eggert, 1977b; Budke et al., 2005); extant plants have either a trilobate and bilobate base morphology. In trilobate species, there are three basal furrows, which run down the lateral face of the corm and join together at the distal end (figure 1a–c). The presence of these basal furrows divide the corm into three sections where secondary growth ultimately results in the formation of the triple corm lobes (Stokey, 1909; Osborn, 1922; Bhambie, 1963; Freund, pers. obs.). In contrast, the bilobate species have only a single furrow, which runs in a line across the base of the corm, dividing it into two halves (figure 1 d–f). While the plants do occasionally acquire additional lobes as they age, the base morphology—the minimum number of lobes the plants have before any elaboration—is consistent (Karrfalt and Eggert, 1977a; b). Also, there are other, rarer morphologies, such as the rhizomatous, mat-forming *I. tegetiformans* Rury (Rury, 1978) and the monolobate *I. andicola* (Amstutz) L.D. Gómez (Amstutz, 1957). However, both *I. andicola* and *I. tegetiformans* begin life with a bilobate morphology before secondarily acquiring these alternate states (Rury, 1978; Karrfalt, 1984b; Tryon et al., 1994); their base morphology is bilobate.

**Fig. 1.**
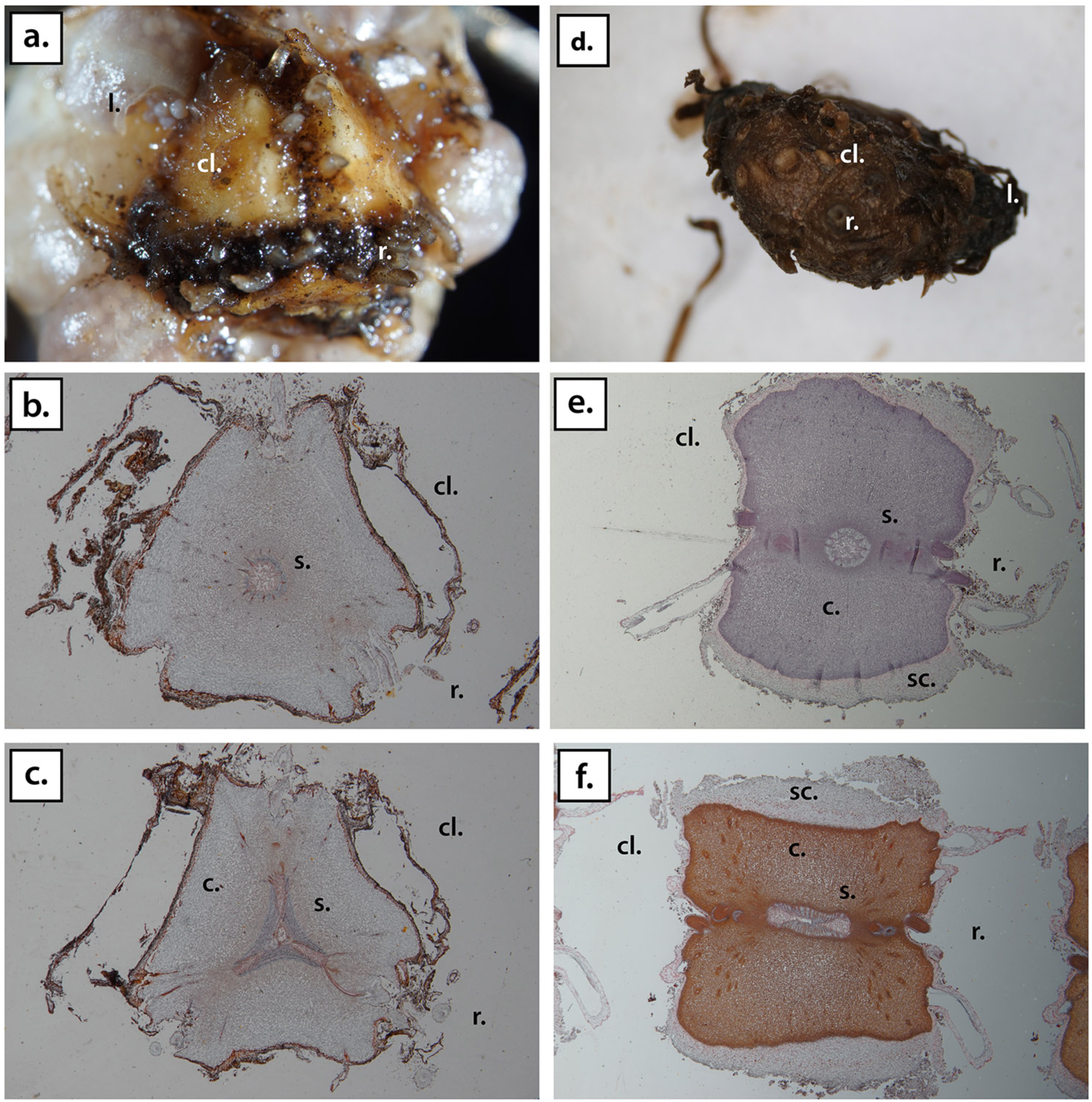
Comparative morphology of tri- and bilobate corms. Trilobed corm – a. *I. nuttallii* (FF266, voucher to be deposited) in proximal view with senesced cortical tissue removed [27.5x, 8.3mm field of view], b.– c. transverse sections of *I. nuttallii* (FF169, RSA796374) below apical rosette (c) and near basal furrows (d) [32.5x, 7.1mm field of view]; Bilobed corm – d. *I. howellii* (FF266.1, voucher to be deposited) in sagittal view [24x, 9.5mm field of view]; e.–f. cross sections of *I. bolanderi* (FF10, RSA811643) below apical rosette (e) and near basal furrow (f) [24.5x, 9.3mm field of view]. Labels: c = cortex ground tissue, cl = corm lobe, r = rootlets, l = leaves, s = stele, sc = senesced cortical ground tissue.

In this study, we explore the evolution of corm lobation across the most recent *Isoëtes* phylogeny (Larsén and Rydin, 2016). If *Isoëtes* has undergone an evolutionary reduction in corm lobation as Karrfelt and Eggart (1977b) hypothesized, we expect the ancestral state for extant *Isoëtes* to be trilobate, and the rate of transition from tri- to bilobate to be higher than the rate of bi- to trilobate. To test this hypothesis, we used reversible-jump Markov chain Monte Carlo (Green, 1995) to explore multiple nonstationary statistical models of morphological evolution (Klopfstein et al., 2015) and to assess whether there is evidence for directional or even irreversible evolution in corm lobation morphology. Finally, we used simulations to examine our power to detect irreversible evolution on datasets of this size.

## METHODS

### Corm lobation characterization

Corm lobe numbers were collected from observations of fresh material, herbarium specimens, and the literature. We characterized lobation from fresh plants for eight species (*Isoëtes appalachiana* D.F. Brunt. & D.M. Britton [four individuals]; *I. bolanderi* Engelm. [20+ individuals]; *I. eatonii* R. Dodge [three individuals]; *I. englemannii* A. Braun [four individuals]; *I. howellii* Engelm. [100+ individuals]; *I. nuttallii* A. Braun ex Engelm. [100+ individuals]; *I. occidentalis* L.F. Hend. [15+ individuals]; and *Isoëtes orcuttii* A.A. Eaton [50+ individuals]). Each individual was cleaned of any encrusting soil, then examined to determine the number of basal furrows and to assess the symmetry of the corm lobes. Plants of different ages were observed for each species when possible to get a sense of variability in lobation pattern as the plants aged, as well as to identify any unusual morphological outliers such as plants with asymmetric lobation or non-linear basal furrows that may have occurred due to either advanced age or damage to the corm. The *I. nuttallii* sampled included two sporelings, which showed distinctly triangular arrangement of their rootlets. Taxa where the plants have a single contiguous furrow and two symmetrical lobes were scored as “bilobate”, while species with three linear furrows with symmetrical lobation around the base of the corm were scored as “trilobate” (Fig. 1).

For lobation numbers garnered from the literature, species with a definitive single reported corm lobation value were assigned that number, while species with a range of reported corm lobe numbers were scored as undefined due to the possibility that the description is of a taxon with a range of corm morphologies, such as *I. tuckermanii* A. Br. (Karrfalt and Eggert, 1977a; Croft, 1980), or due to the inclusion of multiple cryptic species within a single taxon (Table 1).

**Table 1.** Corm lobation numbers for *Isoëtes.* Clade designations follow Larsén and Rydin 2016.

| Species | Observed by<br>F.Freund | Clade | Lobes | Locality | Sources |
| --- | --- | --- | --- | --- | --- |
| *Isoëtes abyssinica*<br>Choiv. – synonym of *I. welwitschii* [sensu (Verdcourt and Beentje, 2002)] |  | B-3 | 3 | Africa, Ethiopia | (Pfeiffer, 1922; Cook, 2004; Crouch et al., 2011) |
| *Isoëtes acadiensis* Kott |  | ? | 2 | USA (MA, ME, NH, NJ, NY, VT),<br>CAN (NB, NF, NS) | (Taylor et al., 1993) |
| *Isoëtes adspersa* A. Braun |  | ? | 3 | Algeria | (Pfeiffer, 1922) |
| *Isoëtes aequinoctialis* A. Braun |  | B-3 | 3 | Angola, South Africa, Mali, Ghana, Tanzania, zambia, Zimbabwe, south to RSA: NAM, NC | (Pfeiffer, 1922; Cook, 2004; Crouch et al., 2011) |
| *Isoëtes alpina* Kirk |  | D-2 | 3 | New Zealand | (Pfeiffer, 1922) |
| *Isoëtes alstonii* C.F. Reed & Verdc. |  | ? | 2 | Egypt; Mozambique; Namibia; Sudan; Tanzania, United Republic of; Zambia; Zimbabwe | (Crouch et al., 2011) |
| *Isoëtes amazonica* A. Braun ex Kuhn |  | A-3 | 3 | Brazil [North (Pará)] | (Pfeiffer, 1922) |
| *Isoëtes anatolica* Prada & Rolleri |  | C | 3 | Turkey | (Prada and Rolleri, 2005) |
| *Isoëtes andicola* (Amstutz) L.D. Gómez | Pers. obs., live plants. | E-2 | 2 | Peru (Lima, Pasco, Junin, Cuzco, Puno), Bolivia | (Karrfalt, 1984b; Tryon et al., 1994) |
| *Isoëtes andina* Spruce ex Hook. – synonym of *I. triquetra* |  | E-2 | 2 | South America, Peru | (Pfeiffer, 1922) |
| *Isoëtes appalachiana* D.F. Brunt. & D.M. Britton | Pers. obs., live plants. | E-2 | 2 | USA (FL, GA, NC, PA, SC, VA) | (Brunton and Britton, 1997) |
| *Isoëtes araucaniana* Macluf & Hickey |  | ? | 2 | Malleco (Chile) | (Macluf and Hickey, 2007) |
| *Isoëtes asiatica* Makino |  | E-2 | 2 | Kamchatka, Sakhalin, the Kuriles and Japan | (Yi and Kato, 2001) |
| *Isoëtes australis* R.O. Williams |  | A-2, D-2 | 2 | Bruce Rock, West Australia | (Williams, 1944) |
| *Isoëtes azorica* Durieu ex Milde |  | ? | 2 | Islands of Azores | (Pfeiffer, 1922) |
| *Isoëtes bolanderi* Engelm. | RSA811637, RSA881638, RSA811639, RSA811643 | E-2 | 2 | USA (AZ, CA, CO, FL, ID, MT, NM, NV, OR, UT, WA, WY), CAN (AB) | (Pfeiffer, 1922; Taylor et al., 1993) |
| *Isoëtes boliviensis* U. Weber |  | ? | 2 | Peru (Cajamarca, San Martin, Ancash, Lima, Pasco, Junin, Ayacucho, Cuzco, Puno), Bolivia (La Paz) | (Tryon et al., 1994) |
| *Isoëtes boryana* Durieu |  | ? | 3 | France (Landes) | (Pfeiffer, 1922) |
| *Isoëtes bradei* Herter |  | A-3 | 3 | Brazil, Sudeste (São Paulo) | Brazilian Flora 2020, (Hickey, 1990) |
| *Isoëtes braunii* Unger |  | ? | 2 | USA (NH, VT, MA), CAN | (Pfeiffer, 1922) |
| *Isoëtes brevicula* E.R.L. Johnson |  | D-2 | 3 | Western Australia | FloraBase |
| *Isoëtes brochoni* Motelay |  | ? | 2 | France | (Pfeiffer, 1922) |
| *Isoëtes butleri* Engelm. |  | ? | 2 | USA (AL, AR, GA, IL, KS, KY, MO, OK, TN, TX) | (Pfeiffer, 1922) |
| *Isoëtes capensis* A.V. Duthie |  | A-1 | 3 | South Africa (Western Cape) | (Cook, 2004; Crouch et al., 2011) |
| *Isoëtes caroli* E.R.L. Johnson |  | D-2 | 3 | Western Australia | FloraBase |
| *Isoëtes caroliniana* (A.A. Eaton) Luebke |  | ? | 2 | USA (NC, AE104TN, VA, WV) | (Taylor et al., 1993) |
| *Isoëtes coreana* Y.H. Chung & H.K. Choi |  | D-3 | 3 | Korea | (Chung, 1986) |
| *Isoëtes coromandeliana* L.f. |  | A-4 | 3 | India | (Pfeiffer, 1922) |
| *Isoëtes cubana* Engelm. ex Baker |  | A-3 | 3 | Cuba (Pinao del Rio), Belize, Mexico (Yucatan) | (Pfeiffer, 1922) |
| *Isoëtes dispora* Hickey |  | ? | 2 | Laguna Tembladera, Lambayeque, Peru | (Tryon et al., 1994) |
| *Isoëtes dixitei* Shende |  | B-3 | 3 | Panchgani, India | (Pant and Srivastava, |
|  |  |  |  |  | 1962) |
| *Isoëtes drummondii* A. Braun |  | D-2 | 3 | Australia (South Australia, Western Australia, Victoria, New South Wales.) | (Osborn, 1922; Pfeiffer, 1922) |
| *Isoëtes duriei* Bory |  | B-1 | 3 | Algeria, Corsica, France, Italy, Turkey | (Pfeiffer, 1922) |
| *Isoëtes eatonii* R. Dodge | Freund - pers. obs. of live plants. | E-2 | 2 | USA (CT, MA, NH, NJ, NY, PA, VT), CAN (ON) | (Pfeiffer, 1922) |
| *Isoëtes echinospora* Durieu |  | E-2 | 2 | CAN (AB, BC, MB, NB, NT, NS, ON, PE, QC, SK, YT), USA (AK, CA, CO, ID, ME, MA, MI, MN, MT, NH, NJ, OH, OR, PA, VT, WA, WI), British Isles, Circum boreal | (Pfeiffer, 1922) |
| *Isoëtes elatior* A. Braun |  | ? | 3 | Tasmania | (Pfeiffer, 1922) |
| *Isoëtes eludens* J.P.Roux, Hopper & Rhian J.Sm. |  | ? | 3 | South Africa (Kamiesberg Mountains, Namaqualand) | (Roux et al., 2009; Crouch et al., 2011) |
| *Isoëtes engelmannii* A. Braun (includes *I. valida*) | Freund - pers. obs. of live plants. | E-2 | 2 | CAN (ON), USA (Al, AR, CT, DE, FL, GA, IL, IN, KY, MD, MA, MI, MO, NH, NJ, NY, NC, OH, PA, RI, SC, TN, VT, VA, WV) | (Pfeiffer, 1922; Taylor et al., 1993) |
| *Isoëtes flaccida* A. Braun |  | E-2 | 2 | USA (Fl, GA) | (Pfeiffer, 1922; Taylor et al., 1993) |
| *Isoëtes flettii* (A.A. Eaton) N. Pfeiff. |  | ? | 2 | USA (WA - Spanaway Lake) | (Pfeiffer, 1922) |
| *Isoëtes foveolata* A.A. Eaton ex R. Dodge |  | ? | 2 | USA (CT, MA, NH), CAN (ONT) | (Pfeiffer, 1922) |
| *Isoëtes gardneriana* Kunze ex Mett. |  | ? | 3 | Brazil [Midwest (Goias)], Paraguay | (Pfeiffer, 1922) |
| *Isoëtes georgiana* Luebke |  | E-2 | 2 | USA (GA) | (Taylor et al., 1993) |
| *Isoëtes giessii* Launert |  | ? | 3 | Namibia (Erongo Mountains & in seasonally wet<br>depressions in<br>Acacia scrub) | (Crouch et al., 2011) |
| *Isoetes gunnii* A. Braun |  | D-2 | 3 | Tasmania (Lake<br>Fenton on Mt.<br>Field) | (Pfeiffer, 1922) |
| *Isoetes habbemensis*<br>Alston |  | D-3 | (2)–3–<br>(4) | New Guinea | (Croft, 1980) |
| *Isoetes hallasanensis*<br>H.K. Choi, Ch. Kim & J.<br>Jung |  | D-3 | 3 | Jeju Island, Korea | (Choi et al., 2008) |
| *Isoetes hawaiiensis* W.C.<br>Taylor & W.H. Wagner |  | E-2 | 2 | Hawaii | (Taylor et al., 1993) |
| *Isoetes heldreichii*<br>Wettst. |  | ? | 3 | Greece (plains of<br>Thessaly, base of<br>Pindus Mts.) | (Pfeiffer, 1922) |
| *Isoetes herzogii* U.<br>Weber |  | E-2 | 2 | Bolivia | (Weber, 1922) |
| *Isoetes hewitsonii*<br>Hickey |  | ? | 2 | Celendin,<br>Cajamarca, Peru | (Tryon et al., 1994) |
| *Isoetes hystrix* Bory &<br>Durieu |  | E-1, E-2 | 3 | Algeria, Italy,<br>France, islands of<br>the Mediterranean | (Pfeiffer, 1922) |
| *Isoetes howellii* Engelm. | RSA796366,<br>RSA796367,<br>RSA796368,<br>RSA796369.<br>RSA796370,<br>RSA796371 | E-2 | 2 | USA (MT, ID, WA,<br>OR, CA) | (Pfeiffer, 1922; Taylor et<br>al., 1993) |
| *Isoetes humilior* A.<br>Braun |  | D-1 | 2 | Tasmania (So.<br>River Esk) | (Pfeiffer, 1922; Taylor et<br>al., 1993) |
| *Isoetes inflata* E.R.L.<br>Johnson |  | ? | 2 | Australia (West<br>Australia) | (FloraBase) |
| *Isoetes japonica* A.<br>Braun |  | D-3 | 3 | Japan (Yokohama) | (Pfeiffer, 1922) |
| *Isoetes jejuensis* H.K.<br>Choi, Ch. Kim & J. Jung |  | D-3 | 3 | Jeju Island, Korea | (Choi et al., 2008) |
| *Isoetes kirkii* A. Braun |  | D-1 | 3 | New Zealand | (Pfeiffer, 1922) |
| *Isoetes labri-draconis*<br>N.R. Crouch |  | ? | 3 | Drakensberg Range<br>in KwaZulu Natal | (Crouch et al., 2011) |
| *Isoetes lacustris* L. |  | E-2 | 2 | CAN (MB, NB,<br>NL, NT, NS, ON,<br>OQ, SK), USA<br>(ME, MA, MI, MN, | (Pfeiffer, 1922; Taylor et<br>al., 1993) |
|  |  |  |  | NH, NY, VT, WI) |  |
| *Isoëtes laosiensis* C. Kim & H.K. Choi |  | A-4 | 3 | Laos | (Kim et al., 2010) |
| *Isoëtes lechleri* Mett. |  | ? | 2 | Argentina, Equador, Colombia, Peru | (Pfeiffer, 1922; Tryon et al., 1994) |
| *Isoëtes lithophila* N. Pfeiff. |  | E-2 | 2 | USA (TX) | (Pfeiffer, 1922; Tryon et al., 1994) |
| *Isoëtes longissima* Bory & Dur. – As *I. velata* forma *longissima* (senu Pfeiffer (1922)) |  | B-3 | 3 | Algeria | (Pfeiffer, 1922) |
| *Isoëtes macrospora* Durieu |  | ? | 2 | Newfoundland to USA (MN) | (Pfeiffer, 1922) |
| *Isoëtes malinverniana* Ces. & De Not. |  | C | 3 | Italy | (Pfeiffer, 1922) |
| *Isoëtes maritima* Underw. |  | E-2 | 2 | Alaska, British Columbia, Washington | (Underwood, 1888) |
| *Isoëtes martii* A. Braun ex Kuhn |  | E-2 | 2 | Brazil [Southeast (Minas Gerais, Rio de Janeiro), South (Rio Grande do Sul)] | Brazilian Flora 2020 |
| *Isoëtes maxima* Hickey, Macluf & Link-Pérez |  | ? | 2 | Brazil [South (Rio Grande do Sul)] | (Hickey et al., 2009) |
| *Isoëtes melanopoda* Gay & Durieu |  | E-2 | 2 | USA (AL, AR, GA, IA, ID, IL, IN, KS, KY, LA, MN, MO, MS, MT, NC, NE, NJ, OK, SC, SD, TN, TX, UT, VA) | (Pfeiffer, 1922; Taylor et al., 1993) |
| *Isoëtes melanospora* Engelm. |  | E-2 | 2 | USA (GA - Stone Mtn.) | (Pfeiffer, 1922; Taylor et al., 1993) |
| *Isoëtes mexicana* Underw. |  | E-2 | 2 | Mexico (Chihuahua, Hidalgo, Mexico, Morelos, Michoacan) | (Pfeiffer, 1922) |
| *Isoëtes mongerensis* E.R.L. Johnson |  | ? | 3 | Western Australia | (FloraBase) |
| *Isoëtes muelleri* A. Braun |  | D-3 | 3 | Eastern Australia (Rockhampton) | (Pfeiffer, 1922) |
| *Isoëtes muricata* Durieu |  | E-2 | 2 | North America | (Pfeiffer, 1922) as *I. braunii* |
| *Isoëtes nigritiana* A. Braun |  | ? | 3 | Nigeria (Along the Niger river, Nupe) | (Pfeiffer, 1922) |
| *Isoëtes novo-granadensis* H.P. Fuchs |  | E-2 | 2 | Columbia, Peru | (Tryon et al., 1994) |
| *Isoëtes nuttallii* A. Braun ex Engelm. | RSA796374, RSA796375, RSA796376 | B-2 | 3 | USA (CA, OR, WA), CAN (Vancouver) | (Pfeiffer, 1922; Taylor et al., 1993) |
| *Isoëtes occidentalis* L.F. Hend. | RSA811640, RSA811641, RSA811642 | E-2 | 2 | USA (CA, WY, CO, ID) | (Pfeiffer, 1922; Taylor et al., 1993) |
| *Isoëtes olympica* A. Braun |  | B-3 | 3 |  | (Pfeiffer, 1922) |
| *Isoëtes orcuttii* A.A. Eaton | Freund - pers. obs. of live plants | B-2 | 3 | USA (CA) | (Pfeiffer, 1922; Taylor et al., 1993) |
| *Isoëtes ovata* N. Pfeiff. |  | ? | 3 | French Guiana, Guyana | (Pfeiffer, 1922) |
| *Isoëtes panamensis* Maxon & C.V. Morton | Freund - pers. obs. at Museo Nacional de Costa Rica | A-3 | 2 | Brazil [Northeast (Maranhão, Bahia), Midwest (Mato Grosso)] | Brazilian Flora 2020 |
| *Isoëtes parvula* Hickey |  | ? | 2 | Laguna Yaurihui, Ayacucho, Perú | (Tryon et al., 1994) |
| *Isoëtes pedersenii* H.P. Fuchs ex E.I. Meza & Macluf |  | ? | 2 | Argentina (Corrientes - Mburucuyá National Park) | (Macluf et al., 2010) |
| *Isoëtes philippinensis* Merr. & R.H. Perry |  | D-3 | 3 | Phillapine islands | (Merrill and Perry, 1940) |
| *Isoëtes piperi* A.A. Eaton |  | ? | 2 | USA (WA) | (Pfeiffer, 1922) |
| *Isoëtes pringlei* Underw. |  | ? | 2 | Mexico (Guadalajara, state of Jalisco) | (Pfeiffer, 1922) |
| *Isoëtes prototypus* D.M. Britton |  | ? | 2 | CAN (N.B., N.S.), USA (MA) | (Taylor et al., 1993) |
| *Isoëtes pseudojaponica* M. Takamiya, Mits. Watan. & K. Ono |  | D-3 | 3 | Japan | (Takamiya et al., 1997) |
| *Isoëtes quiririensis* J.B.S. Pereira & Labiak |  | ? | 2 | Brazil (Serra do Quoriri) | (Pereira and Labiak, 2013) |
| *Isoëtes riparia* Engelm. ex A. Braun |  | ? | 2 | Can (southern region), USA (New England south to DE and PA) | (Pfeiffer, 1922) |
| *Isoëtes saccharata* Engelm. |  | ? | 2 | USA (DE, DC, MD, VA) | (Pfeiffer, 1922) |
| *Isoëtes sampathkumarinii* L.N. |  | D-2 | 2 | India | (Rao, 1944) |
| Rao |  |  |  |  |  |
| *Isoëtes saracochensis*<br>Hickey |  | ? | 2 | Laguna Saracocha,<br>Puno, Peru | (Tryon et al., 1994) |
| *Isoëtes savatieri* Franch. |  | E-2 | 2 | Argentina (Peurto<br>Beuno) | (Hickey et al., 2003) |
| *Isoëtes schweinfurthii*<br>Baker |  | A-4 | 3 | South Africa<br>(Namibia,<br>Botswana,<br>Limpopo), Sudan,<br>Madagascar,<br>Zambia | (Pfeiffer, 1922; Cook,<br>2004; Crouch et al., 2011) |
| *Isoëtes setacea* Lam. |  | E-1 | 3 | France<br>(Montepellier,<br>Herault, Orientals<br>Pyrenees);<br>Morocco; Portugal;<br>Spain (Balears,<br>Spain) | (Pfeiffer, 1922) |
| *Isoëtes stellenbossiensis*<br>A.V. Duthie |  | A-1 | 3 | South Africa<br>(western part of<br>Western Cape<br>province) | (Cook, 2004; Crouch et al.,<br>2011) |
| *Isoëtes stephansenii*<br>A.V. Duthie |  | A-1 | 2 | South Africa (flasts<br>around Stellenbosch<br>in the Western<br>Cape) | (Crouch et al., 2011) |
| *Isoëtes stevensii* J.R.<br>Croft |  | D-3 | (2)–3–<br>(4) | New Guinea | (Croft, 1980) |
| *I. storkii* T.C. Palmer | GH00021453,<br>CM0102,<br>NY00144272<br>(virtual<br>herbarium<br>sheets),<br>Freund - Pers.<br>obs. at CR | E-2 | 2 | Costa Rica | JStor Global Plants virtual<br>herbarium<br>http://plants.jstor.org/search?filter=name&so=ps_group_by_genus_species+asc&Query=isoetes+storkii,<br>accessed 12/28/2016 |
| *Isoëtes subinermis* Cesca<br>– synonym of *I. histrix* |  | B-1 | 3 |  | (Cesca and Peruzzi, 2001) |
| *Isoëtes taiwanensis* De<br>Vol | Freund - Pers.<br>obs. of live<br>plants | D-3 | 3 | Taiwan | (Chiang, 1976) |
| *Isoëtes tegetiformans*<br>Rury |  | ? | 2 | USA (GA) | (Rury, 1978; Taylor et al.,<br>1993) |
| *Isoëtes toximontana*<br>Musselman & J.P. Roux |  | A-1 | 3 | South Africa<br>(Giftberg endemic) | (Cook, 2004; Crouch et al.,<br>2011) |
| *Isoëtes transvaalensis*<br>Jermy & Schelpe |  | ? | 3 | Lesotho, South<br>Africa (Limpopo,<br>Mpumalanga, Free | (Cook, 2004; Crouch et al.,<br>2011) |
|  |  |  |  | State, KwaZulu-Natal) |  |
| *Isoëtes triquetra* A. Braun |  | E-2 | 2 | Peru | (Pfeiffer, 1922) |
| *Isoëtes truncata* (A.A. Eaton) Clute |  | ? | 2 | CAN (Vancouver Island), USA (Alaska) | (Pfeiffer, 1922) |
| *Isoëtes tuerckheimii* Brause |  | E-2 | 2 | Dominican Republic (Santo Domingo) | (Pfeiffer, 1922) |
| *Isoëtes valida* (Engelm.) Clute |  | E-2 | 2 | USA (N.C., Tenn, Va., W. VA.) | (Taylor et al., 1993) |
| *Isoëtes velata* A. Braun |  | B-3 | 3 | Italy (Algeria, Corsica, Sicily) | (Pfeiffer, 1922) |
| *Isoëtes welwitschii* A. Braun ex Kuhn |  | ? | 3 | Angola, South Africa (Mpumalanga, KwaZulu-Natal), Madagascar | (Pfeiffer, 1922; Cook, 2004; Crouch et al., 2011) |
| *Isoëtes wormaldii* R. Sim |  | ? | 3 | South Africa (Eastern Cape - small area between Grahamstown & East London) | (Pfeiffer, 1922; Cook, 2004; Crouch et al., 2011) |
| *Isoëtes yunguiensis* Q.F. Wang & W.C. Taylor |  | D-3 | 3 | China | (Qing-Feng et al., 2002) |

Additionally, three accessions were coded as unknown for lobation due to identification uncertainties: 1) One accession of *I. savatieri* Franchet was collected in Uruguay, outside of the currently accepted range of the species (Hickey et al., 2003), and we were unable to examine the specimen to determine its identity or lobation. 2) *I. histrix* Bory & Durieu is a known species complex (Bagella et al., 2011; 2015) and specimens fall in two different areas of our phylogeny. One is sister to *I. setacea* Lam., a position that is consistent with morphology and with existing hypotheses of their relationship (Hoot et al., 2006; Bagella et al., 2011; Troia and Greuter, 2014); we treated this accession as correctly identified. The second accession falls phylogenetically distant from the first, and presumably is misidentified; we were unable to ascertain its true identity or morphology, so coded it as unknown. 3) *I. australis* R.O. Williams also shows up in two places in the phylogeny: Clade A and Clade D. The Clade A plant was collected and identified by Dr. Carl Taylor, an authority on *Isoëtes*, so we treated this accession as correctly identified and coded the clade D plant as unknown. All character states and literature sources can be found in Table 1.

### Phylogenetic modeling

Ancestral character state reconstructions were performed on a posterior sample of 15000 trees from Larsén and Rydin (2016) that we rooted on the bipartition between Clade A and the remainder of the genus (following Larsén and Rydin 2016).

We employed reversible-jump MCMC (Green, 1995) in RevBayes (Höhna et al., 2016) to explore the space of all five possible continuous-time Markov models of phenotypic character evolution and to infer ancestral states. The reversible-jump MCMC sampled from the five models in proportion to their posterior probability. This approach enabled model fit comparisons through Bayes factors (Kass and Raftery, 1995), and provided the opportunity to account for model uncertainty by making model-averaged ancestral state and parameter estimates (Madigan and Raftery, 1994; Kass and Raftery, 1995; Huelsenbeck et al., 2004; Freyman and Hoehna, 2017). The five models of corm lobation evolution considered were: a model with the rate of lobation gain and loss set to be equal (the 1-rate model); a model where the rates of lobation gain and loss are independent and non-zero (the 2-rate model); two irreversible models where the rate of either lobation gain or loss was fixed to zero; and lastly a model where both rates were fixed to zero. To test for directional evolution we used nonstationary models of character evolution with root state frequencies that differed from the stationary frequencies of the process (Klopfstein et al., 2015).

Each of the five models was assigned an equal prior probability using a uniform set partitioning prior. The root state frequencies were estimated using a flat Dirichlet prior. The rates of corm lobation gain and loss were drawn from an exponential distribution with a mean of 1 expected character state transitions over the tree (λ=τ/1where τ is the length of tree).

The MCMC was run for 22000 iterations, where each iteration consisted of 48 MCMC proposals. The 48 proposals were scheduled randomly from six different Metropolis-Hastings moves that updated the sampled tree, root frequencies, and corm lobation gain and loss rate parameters. The first 2000 iterations were discarded as burnin, and samples were logged every 10 iterations. Convergence of the MCMC was confirmed by ensuring that the effective sample size of all parameters was over 600. The results were summarized and plotted using the RevGadgets R package (https://github.com/revbayes/RevGadgets). The scripts that specify our model, run the analysis, and summarize results are available in the code repository at https://github.com/wf8/isoetes.

### Simulations

To test how many observed characters are necessary to reliably infer irreversible evolution on a phylogeny the size of ours, we simulated 10 datasets with each of 1, 5, 10, 50, or 100 characters per tip (for a total of 50 simulations; note, our empirical dataset has a single character per tip). Each dataset was simulated under an irreversible model with the mean rate of corm lobation loss set to the value estimated by the irreversible model using the observed corm lobation data (2.39 changes per unit branch length). We performed the simulations using RevBayes over the maximum a posteriori phylogeny from the same tree distribution used to infer the ancestral states. For each of the 50 simulated datasets an MCMC analysis was run for 11000 iterations, with the first 1000 iterations dropped as burnin. The model used was identical to that used for the observed corm lobation dataset, except that for the simulated datasets we fixed the maximum a posteriori phylogeny instead of integrating over the posterior distribution of trees.

## RESULTS

### Model fit comparisons

The maximum a posteriori model of corm lobation evolution was the tri- to bi- irreversible model (which did not allow transitions from the bilobate to the trilobate state) with a posterior probability of 0.38 (Table 2). This tri- to bi- irreversible model was weakly supported over the 1-rate and 2-rate reversible models (Bayes factor = 1.26 and 1.21, respectively; Kass and Raftery, 1995); however, all three models were strongly supported over the bi- to tri- irreversible model. Since the Bayes factor support for best-supported model over the next two was negligible, we focus mostly on the model-averaged parameter estimates and ancestral states.

**Table 2.** Comparisons of models of corm lobation evolution.

|  |  | Bayes factors |  |  |  |
| --- | --- | --- | --- | --- | --- |
|  | model posterior probability | bi- to tri- irreversible | tri- to bi- irreversible | 1-rate | 2-rate |
| bi- to tri- irreversible | 0.0 | - | <1 | <1 | <1 |
| tri- to bi- irreversible | 0.38 | >1000 | - | 1.26 | 1.21 |
| 1-rate | 0.30 | >1000 | <1 | - | <1 |
| 2-rate | 0.32 | >1000 | <1 | 1.04 | - |

### Model averaged parameter estimates and ancestral states

The model-averaged estimated rate of transition from tri- to bilobate forms was significantly non-zero (mean = 2.17 changes per unit branch length, 95% HPD interval: 0.015–5.69), whereas the rate of bi- to trilobate transitions was not significantly non-zero (mean = 0.82, 95% HPD interval: 0.0–3.35; Fig. 2). The model-averaged maximum a posteriori ancestral state of *Isoëtes* was trilobate with a posterior probability of 1.0 (Fig. 3). The ancestral state of the New World clade (Fig. 3, “Clade E-2”) was bilobate with a posterior probability of 0.99. The bilobate morphology arose independently in six places over the phylogeny (Fig. 3). No reversals from bilobate to trilobate were inferred. All species with unknown base corm lobation characters (*I. stevensii* J.R. Croft*, I. habbemensis* Alston, and *I. hallasanensis* H. K. Choi, Ch. Kim & J. Jung) were derived from a trilobate most recent common ancestor, with a posterior probability near 1.0.

**Fig. 2.**
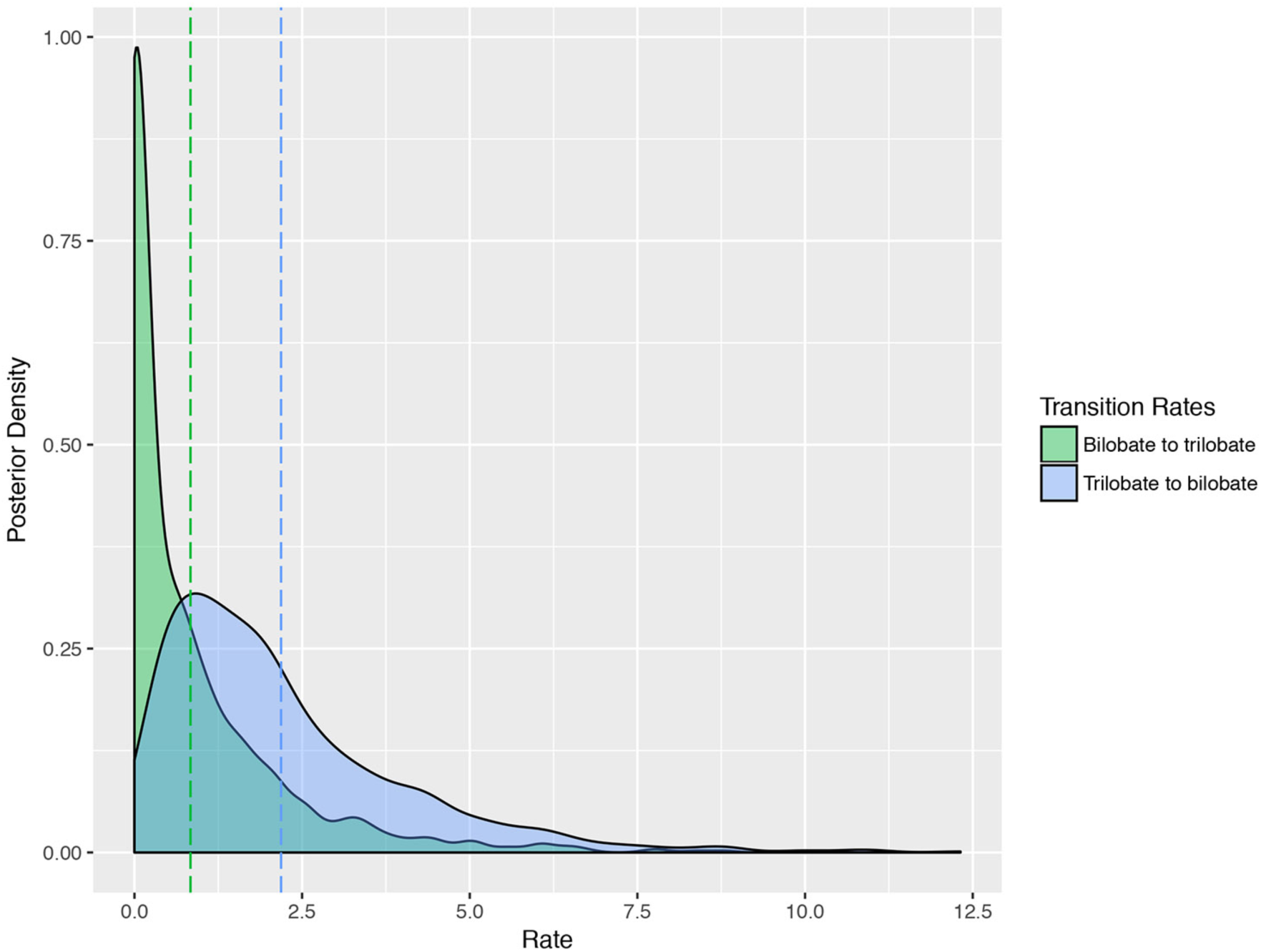
Model-averaged posterior densities of transition rates between corm lobation states. Mean values are represented by dashed lines. The rate of transition from tri- to bilobate forms was significantly non-zero (mean = 2.17, 95% HPD interval: 0.015–5.69), whereas the rate of bi- to trilobate transitions was not significantly non-zero (mean = 0.828.00, 95% HPD interval: 0.0–3.3526.08). Transition rates are reported in changes per unit branch length.

**Fig. 3.**
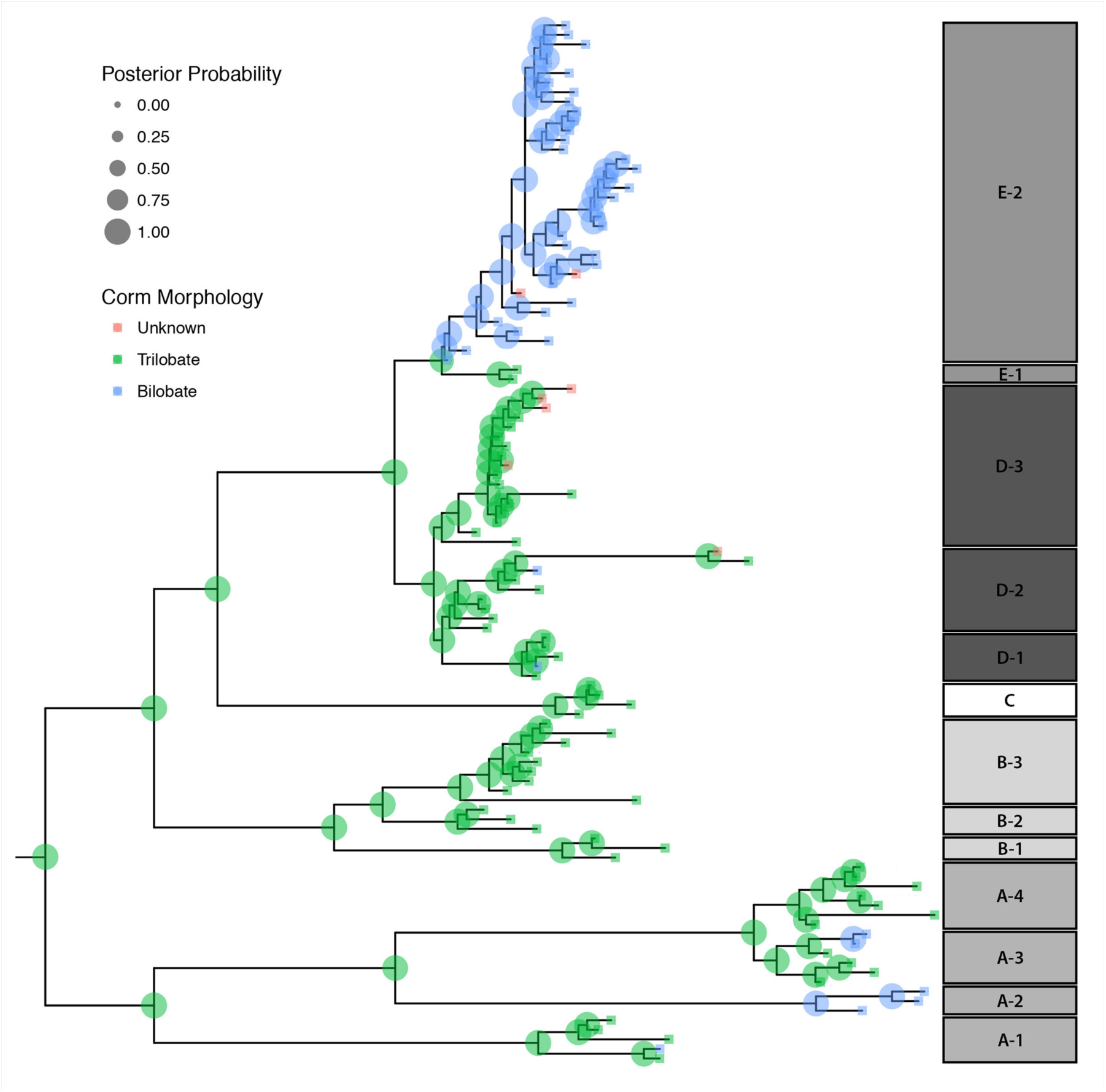
Bayesian model-averaged ancestral states of *Isoëtes* corm lobation inferred over the posterior tree sample from Larsén and Rydin (2016). Ancestral states are summarized on the maximum a posteriori phylogenetic tree. The size of the circles at each node represent the posterior probability of the most probable ancestral state, and the color represents the state: green = trilobate corms, blue = bilobate corms, and red = unknown. Boxes to the right of the figure reflect subclade designations to broad subclade: A = Gondwanan Clade, B = Laurasian Clade, C = Italian Clade, D = Austro-Asian clade, and E = the New World or American Clade. See Table 1 for specific placement of taxa into sub-subclades.

### Simulations

For simulated datasets with a single character the true irreversible model was at best weakly supported over the other models; the mean posterior probability of the true irreversible model was 0.37 (range = 0.22–0.44). As the number of characters increased, the posterior probability of the true model increased (Fig. 4). With 100 characters the support for the true model was strong; the mean posterior probability was 0.85 (range = 0.80–0.92).

**Fig. 4.**
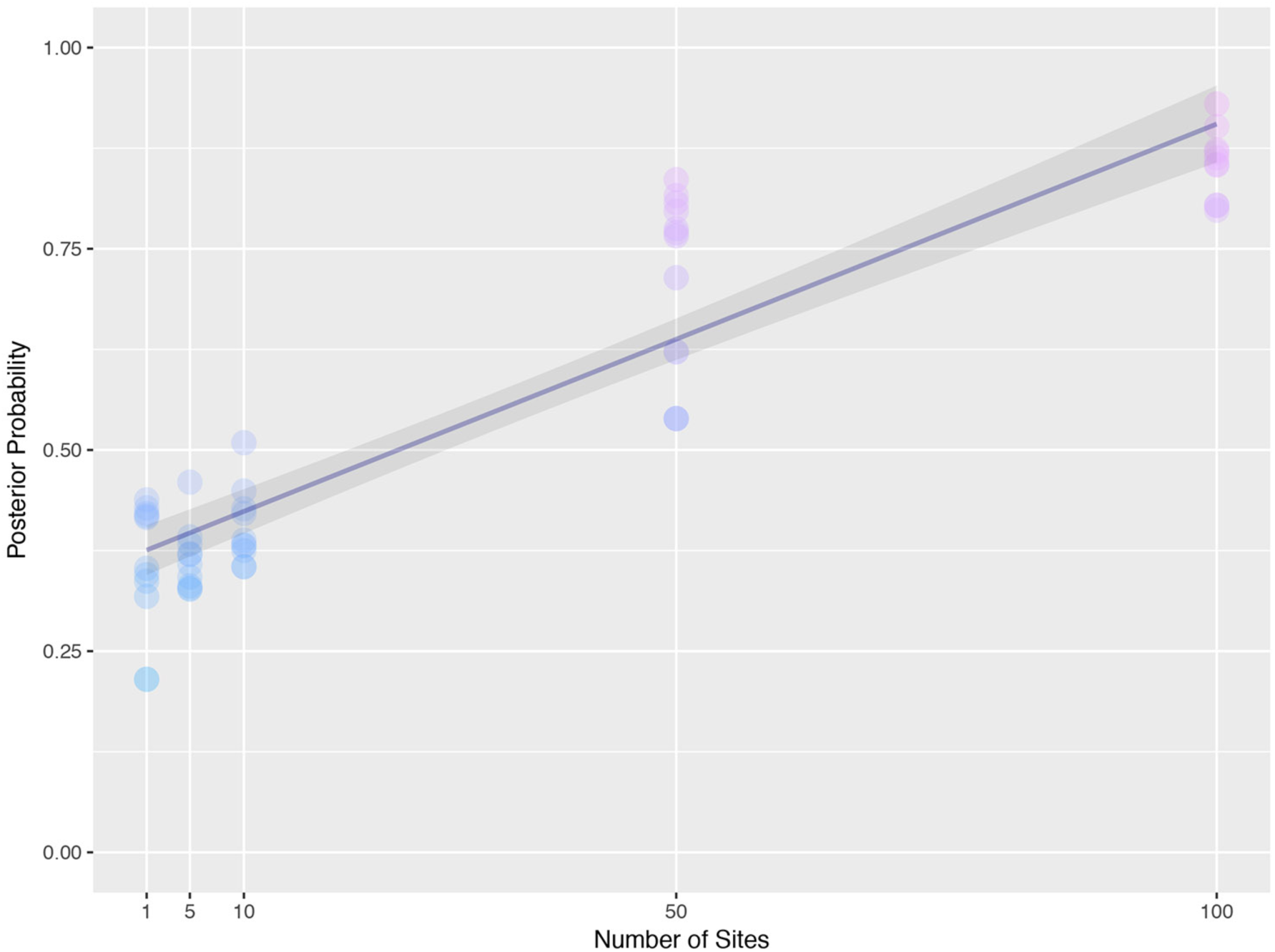
The statistical power to detect irreversible evolution as a function of the number of characters available. Each point plotted represents a different simulation replicate. The y-axis shows the posterior probability of the true irreversible model of character evolution. The x-axis shows the number of simulated characters (sites or columns in the data matrix). Ten replicates were simulated for 1, 5, 10, 50, and 100 character datasets, resulting in a total of 50 simulated datasets.

## DISCUSSION

Our analyses support the hypothesis of directional reduction in lobe number over time in *Isoëtes*, with the best-supported model being one of irreversible evolutionary reduction. Additionally, even when incorporating model and phylogenetic uncertainty and allowing for reversals, the model-averaged estimate of the transition rate from tri- to bilobate was much higher than the estimate of the transition rate from bi- to trilobate (the latter rate was not significantly non-zero). Furthermore, we found strong support for the ancestral state for all extant *Isoëtes* being trilobate, indicating that there have been multiple convergent reductions of the corms to bilobate. These bilobate forms are nested deeply within clades of trilobate plants. These results support the hypothesis that crown *Isoëtes* has continued a reduction in corm morphology from the larger arborescent lycophytes. Additionally, while the bilobate form has emerged multiple times throughout the phylogeny, it is the dominant morphology in only one major clade: the “American clade” (Clade E-2, sensu Larsén and Rydin 2016; Fig. 3). In this clade, nearly all species have a bilobate morphology.

The simulations demonstrate that the relatively weak support for the irreversible reductionary model over the reversible models is likely due to the inherent limitation in statistical power of a single observed morphological character over a phylogeny of this size. Repeating this analysis with a larger, more densely sampled phylogeny, or one that incorporates fossil data, might find stronger support for the irreversible model of corm morphology evolution. Nevertheless, it is in cases like this where no single model is decisively supported over others that reversible-jump MCMC and Bayesian model-averaging demonstrate their utility for testing phylogenetic hypotheses of character evolution (Huelsenbeck et al., 2004; Freyman and Hoehna, 2017).

While the bilobate morphology dominates only Clade E-2, it does occur in several other areas of the phylogeny (Fig. 2, Table 1). In areas where Clade E-2 co-occurs with others, such as South America, and the west coast of the United States, corm lobation is in determining which clade a plant belongs to. However, outside the range of Clade E-2, other bilobate taxa occur as single species nested within larger trilobate clades. As such, using corm lobation outside the Americas to assign plants with unknown phylogenetic placement is not advisable, since they may represent other independent evolutions of the character state.

When assessing corm lobation numbers, it is important to determine if the corm lobe numbers are the base lobe numbers, or additional lobes that have developed as the plant ages (Stokey, 1909; Karrfalt and Eggert, 1977a; b; Freund, pers obs.). While Karrfalt and Eggert (1977b) reported a propensity for gaining additional lobes in their study of *I. tuckermanii* (68% bilobate, 30% trilobate, 2% tetralobate), other observers, working on other taxa, have not found this degree of variability (Engelmann, 1882; Freund pers. obs. of over 200 specimens of *I. howellii* and 400 specimens of *I. nuttallii*). In fact, we have observed fewer than ten total *I. howellii* and *I. nuttallii* specimens that were not either bilobate or trilobate, respectively (Freund, pers obs). The degree of variability in *I. tuckermanii* and the stability of *I. howellii* and *I. nuttallii* show that when describing corm lobation numbers, sampling multiple individuals to assess the degree of variability within a given species, and to determine the base number of lobes, is critical. If at all possible, it is worthwhile to look at very young plants, especially sporelings, to determine the base lobe number.

## CONCLUSIONS

Our results support the hypothesis that crown *Isoëtes* have continued an evolutionary reduction in corm morphology from the larger arborescent lycophytes, with the best-supported model being one of irreversible evolutionary reduction. However, results from our simulation study showed that a dataset of this size only has weak statistical power to support irreversible models of character evolution, emphasizing the need for broader sampling of both extant and fossil taxa. When we accounted for the uncertainty in character evolution models by making model-averaged estimates, we found strong support for the hypothesis of directional evolutionary reduction in corm number, with the rate of lobe loss estimated to be much higher than the rate of lobe gain. Furthermore, we found strong support that the ancestral state for all extant *Isoëtes* was trilobate, indicating that there have been multiple convergent reductions of the corms to bilobate.

## ACKNOWLEDGEMENTS

The authors thank Eva Larsén and Catarina Rydin for providing the phylogenies used in this analysis, the U.C. Berkeley Herbaria and Department of Integrative Biology, the Rancho Santa Anna Botanic Garden and Herbarium, and the National Herbarium of Costa Rica for use of their collections to confirm corm lobation numbers, and the National Forest Service, Bureau of Land Management, California Department of Fish and Wildlife, and Sonoma Land Trust for allowing us to collect specimens on their properties.

## APPENDICIES

## Appendix 1

Vouchers examined for morphological determinations for this study. *I. bolanderi*: RSA811637, RSA881638, RSA811639, RSA811643. *I. howellii*: RSA796366, RSA796367, RSA796368, RSA796369, RSA796370, RSA796371, UC (tbd). *I. nuttallii* RSA796374, RSA796375, RSA796376, UC (tbd). *I. occidentalis*: RSA811640, RSA811641, RSA811642. *I. orcuttii* UC (tbd). *I. storkii*: GH00021453, CM0102, NY00144272 (virtual herbarium sheets)

